# Simultaneous recording of spikes and calcium signals in odor-evoked responses of *Drosophila* antennal neurons

**DOI:** 10.1101/2025.06.27.662059

**Authors:** Yiyi Xiao, Shiuan-Tze Wu, Yinan Xuan, Scott A Rifkin, Chih-Ying Su

**Affiliations:** Department of Neurobiology, University of California San Diego, La Jolla, CA 92093, USA; Department of Electrical and Computer Engineering, University of California San Diego, La Jolla, CA 92093, USA; Department of Ecology, Behavior, and Evolution, University of California San Diego, La Jolla, CA 92093, USA; Division of Biology and Biological Engineering, California Institute of Technology, Pasadena, CA 91125, USA

**Author notes:** Corresponding author: Chih-Ying Su.

**Keywords:** Olfactory receptor neurons, *D. melanogaster*, ab2A, Or59b, dual-modality recording, single-sensillum recording, widefield calcium imaging, GCaMP, antenna

## Abstract

Most insects, including agricultural pests and disease vectors, rely on olfaction for key innate behaviors. Consequently, there is growing interest in studying insect olfaction to gain insights into odor-driven behavior and to support efforts in vector control. Calcium imaging using GCaMP fluorescence is widely used to identify olfactory receptor neurons (ORNs) responsive to ethologically relevant odors. However, accurate interpretation of GCaMP signals in the antenna requires understanding both response uniformity within an ORN population and how calcium signals relate to spike activity. To address this, we optimized a dual-modality recording method combining single-sensillum electrophysiology and widefield imaging for *Drosophila* ORNs. Calcium imaging showed that homotypic ab2A neurons exhibit similar odor sensitivity, consistent with spike recordings, indicating that a single ORN’s response can reliably represent its homotypic counterparts. Furthermore, concurrent dual recordings revealed that peak calcium responses are linearly correlated with spike activity, regardless of imaging site (soma or dendrites), GCaMP variant, odorant, or fly age. These findings validate the use of somatic calcium signals as a reliable proxy for spike activity in fly ORNs and provide a foundation for future large-scale surveys of spike–calcium response relationships across diverse ORN types.

## Introduction

Most insect species, including agricultural pests and disease vectors, rely on olfaction to guide critical innate behaviors such as host-seeking, foraging, courtship, and egg-laying (Afify et al., 2019; Cao et al., 2020; Chang et al., 2017; DeGennaro et al., 2013; Dormont et al., 2021; Fandino et al., 2019; Guo et al., 2020; Kim et al., 2017; Konopka et al., 2021; Lahondère et al., 2020; Liu et al., 2020; Manwill et al., 2020; McBride et al., 2014; McMeniman et al., 2014; Raji et al., 2019; Wheelwright et al., 2021). As a result, there is growing interest in insect olfaction in order to understand odor-guided behavior and also inform vector control strategies.

With the growing use of advanced transgenic and genome editing technologies, *in vivo* calcium imaging has emerged as a key method for identifying olfactory receptor neurons (ORNs) that respond to host-related or other ethologically relevant odors across diverse insect species (Afify et al., 2019; Bui et al., 2019; Carcaud et al., 2023; Fujiwara et al., 2014; Hart et al., 2023; Jiang et al., 2024; Lahondère et al., 2020; Mariette et al., 2023; Matthews et al., 2019; Wang et al., 2003; Zhao et al., 2022). Unlike electrophysiological recordings, calcium imaging can simultaneously monitor odor-evoked activity across genetically defined neuronal populations. And unlike other genetically encoded calcium indicators such as CaLexA (Masuyama et al., 2012), TRIC (Gao et al., 2015), CaMPARI (Fosque et al., 2015), or CRTC (Bonheur et al., 2023), GCaMP-based live imaging enables direct quantification of dose–response relationships in the same neurons, allowing for accurate measurement of odor sensitivity.

To measure an ORN’s calcium activity, responses can be recorded either peripherally in the antenna via transcuticular imaging or centrally at the presynaptic axon terminals in the antennal lobe. While antennal lobe imaging is well-established in *Drosophila* (Wang et al., 2003) and has been successfully applied in other insect species (Carcaud et al., 2023; Fujiwara et al., 2014; Hart et al., 2023; Jiang et al., 2024; Lahondère et al., 2020; Melo et al., 2019; Vinauger et al., 2018; Wolff et al., 2023; Zhao et al., 2022), its broader application is constrained by the advanced technical requirements, namely, the delicate removal of head cuticle and access to specialized multiphoton microscopy. Additionally, axons from multiple ORN types can converge on a single glomerulus, making it difficult to resolve activity from individual neuronal populations. For instance, two ORN subsets expressing the Amt receptor—one in the ac1 sensilla and the other in the sacculus—both project to the same glomerulus. Nevertheless, widefield calcium imaging of the antenna allowed us to distinguish their ammonia responses based on their distinct antennal locations, revealing differences in their calcium response dynamic ranges (Vulpe et al., 2021).

These results further suggest that an ORN’s somatic calcium response is influenced not only by its receptor identity but potentially also by other factors, such as the composition of calcium channels, transporters, buffering proteins, and the neuron’s intrinsic electrotonic properties (Ali & Kwan, 2019; Friel & Chiel, 2008; Helmchen & Tank, 2015; Huang et al., 2021). For example, a 10% increase in fluorescence signal may reflect a high-frequency, near-saturating spike response in one ORN type (Vulpe et al., 2021), but represent low responses in another on the rising phase of dosage curve (Verschut et al., 2023). Therefore, to accurately interpret antennal calcium imaging data, particularly when comparing across ORN types, it is critical to understand how calcium signals correlate with the underlying ground truth of neuronal activity: spike frequency.

This information is especially important given the remarkable diversity of insect ORNs, which are housed in sensory hairs named sensilla that are categorized into distinct morphological classes (Ng et al., 2020). Correspondingly, ORNs from different sensillum classes also exhibit characteristic morphological features. For example in *D. melanogaster*, the outer dendrites of basiconic and intermediate ORNs are numerously branched, whereas those of coeloconic and trichoid neurons are typically unbranched (Nava Gonzales et al., 2021; Shanbhag et al., 1999). Our previous serial block-face scanning electron microscopy study further shows that these ORN classes differ significantly in dendritic length and overall size (Nava Gonzales et al., 2021), which in turn contribute to their differing electrotonic properties (Zhang et al., 2019).

To determine whether the relationship between odor-evoked spike activity and calcium signals varies across ORN types, it is essential to first establish a method for simultaneously measuring both responses, and to address key factors that may influence calcium signal analysis. These are the central objectives of this study.

## Methods

### Animals

All *Drosophila melanogaster* were reared on standard cornmeal-molasses food at 25 °C with ∼60% humidity under a 12-hour light/dark cycle. Flies were collected upon eclosion, separated by sex, and housed in groups of approximately 10. They were transferred to fresh food vials every other day. Unless otherwise specified, all experimental flies were 4-day-old virgin females. Transgenic fly lines used in this study were obtained from the Bloomington *Drosophila* Stock Center. GCaMP6m (RRID:BDSC_42750), GCaMP7c (RRID:BDSC_80908), and GCaMP8s (RRID:BDSC_92593) were expressed in ab2A ORNs using the *Or59b-GAL4* driver (RRID:BDSC_23897). The full genotypes were as follows:

*w^-^; Or59b-GAL4/+; 20xUAS-IVS-GCaMP6m in VK00005/+*
*w^-^; Or59b-GAL4/20xUAS-IVS-jGCaMP7c in su(Hw)attP5; +*
*w^-^; Or59b-GAL4/+; 20xUAS-IVS-jGCaMP8s in VK00005/+*

### Widefield antennal calcium imaging

To prepare for antennal calcium imaging, the head was inserted into the narrow end of a truncated 200-μl plastic pipette tip, leaving the antenna exposed. The antenna was then stabilized between a tapered glass microcapillary and a coverslip coated with double-sided type.

Images were acquired via Micro-Manager 1.4 (Edelstein et al., 2014) at a resolution of 706 × 706 pixels (155.32 × 155.32 µm) with a 2 × 2 binning. Imaging was performed with a back-illuminated scientific CMOS camera (Prime 95B, Photometrics) mounted on an upright microscope (Olympus BX51WI) equipped with a 50× air objective (NA 0.50, LMPlanFl, Olympus). GCaMPs were excited using 50-ms blue light pulses (470 nm) delivered at 50% duty cycle and 80% of full LED power (CoolLED pE-4000, Universal LED Illumination System). Image acquisition was conducted at 10 Hz for a total of 380 frames (∼38 seconds), including a 7-second pre-stimulus baseline.

Motion correction was performed with the “moco” algorithm in ImageJ (Dubbs et al., 2016). For each antenna, the “algorithm-selected” region of interest (ROI) was determined as the sensillum or soma showing the highest fluorescence change (ΔF/F_0_) in response to the second-highest odorant concentration. The same ROI was then applied to all images acquired from the same antenna at other odorant concentrations.

A custom MATLAB script was used to calculate odor-induced changes in calcium fluorescence (ΔF/(F_0_ − F_bac_), referred to as ΔF/F for simplicity). ΔF was defined as F_signal_ − F_0_, where F_signal_ is the instantaneous fluorescence pixel value averaged across the ROI, and F_0_ is the mean fluorescence of the same ROI during the 2-second pre-stimulus period. To correct for variable background fluorescence, a region of the same size as the ROI was manually selected from a dark area near the edge of the antenna. The average pixel value from this background region (F_bac_) during the 2-second pre-stimulus period was subtracted from F_0_ to determine the baseline calcium fluorescence. Odor-evoked calcium fluorescence traces were smoothed in MATLAB using a sliding window (window length: 5; overlap: 4). Pseudocolored heatmaps of peak calcium responses (ΔF) were generated and further processed using a Gaussian bilateral filter in MATLAB.

### Electrophysiology

ab2 sensilla were identified based on their characteristic positions on the antenna, A/B spike amplitude ratio, and their odor response profiles (Grabe et al., 2016; Hallem & Carlson, 2006; Nava Gonzales et al., 2021; Zhang et al., 2019). Single-sensillum recording was performed as follows. Briefly, the electrical activity of the target ORN was recorded extracellularly by inserting a sharp glass electrode filled with adult hemolymph-like solution (AHL) (Stewart et al., 1994) into the sensillum. A reference electrode, also filled with AHL, was inserted in the eye. Data acquisition was controlled using Clampex 10.7 (Molecular Devices). AC signals (100– 20,000 Hz) were recorded and amplified using an NPI EXT-02F amplifier (ALA Scientific Instruments) and digitized at 5 kHz with Digidata 1550 (Molecular Devices). Each recording session was for 10 seconds, including a 1-second pre-stimulus baseline. Six ab2 sensilla were recorded per antenna per fly.

ORN spikes were sorted using a custom MATLAB routine (Martelli & Fiala, 2019). Peri-stimulus time histograms (PSTHs) were generated by averaging spike activity in 50-ms bins and smoothed using MATLAB’s sliding window function (window length: 5; overlap: 4). For adjusted PSTHs, the baseline spike rate—calculated from a 1-second pre-stimulus period—was subtracted from each binned value.

### Dual-modality recording

For simultaneous calcium imaging and single-sensillum recording, widefield imaging was performed as described above. For concurrent single-sensillum recording, tungsten electrodes were used because they provided greater recording stability than glass electrodes during the longer recording duration, about 10 minutes per antenna. Tungsten electrodes were freshly prepared prior to each experiment. Specifically, a tungsten rod (0.01 × 3 inch, 717000, A-M Systems) mounted in an electrode holder (ST50-BNC, Siskiyou) was electrochemically sharpened in 0.5 N KOH using a microelectrode etcher (EE-1D, Bak Electronics) at 20V for nine to ten cycles.

AC signals (100–20,000 Hz) were recorded and amplified using an NPI EXT-02F amplifier (ALA Scientific Instruments) and digitized at 5 kHz with Digidata 1550 (Molecular Devices). Data acquisition was controlled using Clampex 10.7 (Molecular Devices), which also triggered image capture in Micro-Manager to enable synchronized calcium imaging.

### Odorants

Methyl acetate (Sigma 45999) and ethyl acetate (Sigma 270989) were used to activate ab2A neurons. Lower concentrations were prepared by serial volume-to-volume dilutions in paraffin oil (Sigma 76235) from stock solutions. For each stimulus, 100 µL of odorant was applied to a filter paper disc placed inside a glass Pasteur pipette. Odor pulses were delivered at 300 mL/min for 300 ms into a main humidified airstream flowing at 2 L/min.

### Fitting and statistical analysis

The Hill equation was fitted using MATLAB’s “lsqcurvefit” function to determine EC_50_ values. The same function was also used to estimate response kinetics by modeling the rise and decay phases with single exponential functions, from which the time to reach 63% of the peak (τ_on_) and to decay to 37% of the peak (τ_off_) were calculated. Linear regressions were performed using MATLAB’s “polyfit” function.

For comparisons of pEC_50_ values—defined as –logEC_50_—or dendritic calcium responses across flies, overall significance was assessed using one-way Analysis of Variance (ANOVA) (Figures 1C, 1F, and 2C). For comparisons of baseline calcium fluorescence across ages, one-way ANOVA followed by the Tukey–Kramer test was used to assess pairwise *P* values (Figure 4A). Comparisons of peak dendritic calcium responses and pEC_50_ values between SSR-recorded and algorithm-selected ROIs were performed using the two-sided Wilcoxon rank-sum test (Figure 2D). For comparisons of dose–response curves between somatic and dendritic calcium responses, two-way analysis of covariance (ANCOVA) was applied (Figure 2G). Linear relationships between spike and calcium responses were evaluated using Pearson correlation to obtain R and *P* values. To compare spike–calcium relationships across multiple groups, one-way multivariate analysis of variance (MANOVA) with Pillai’s Trace was used (Figure 3), followed by the Tukey–Kramer test to assess pairwise *P* values when there were more than two groups (Figures 2H and 4B). Further details on sample sizes and statistical significance for each test are provided in the figure legends.

**Figure 1.**
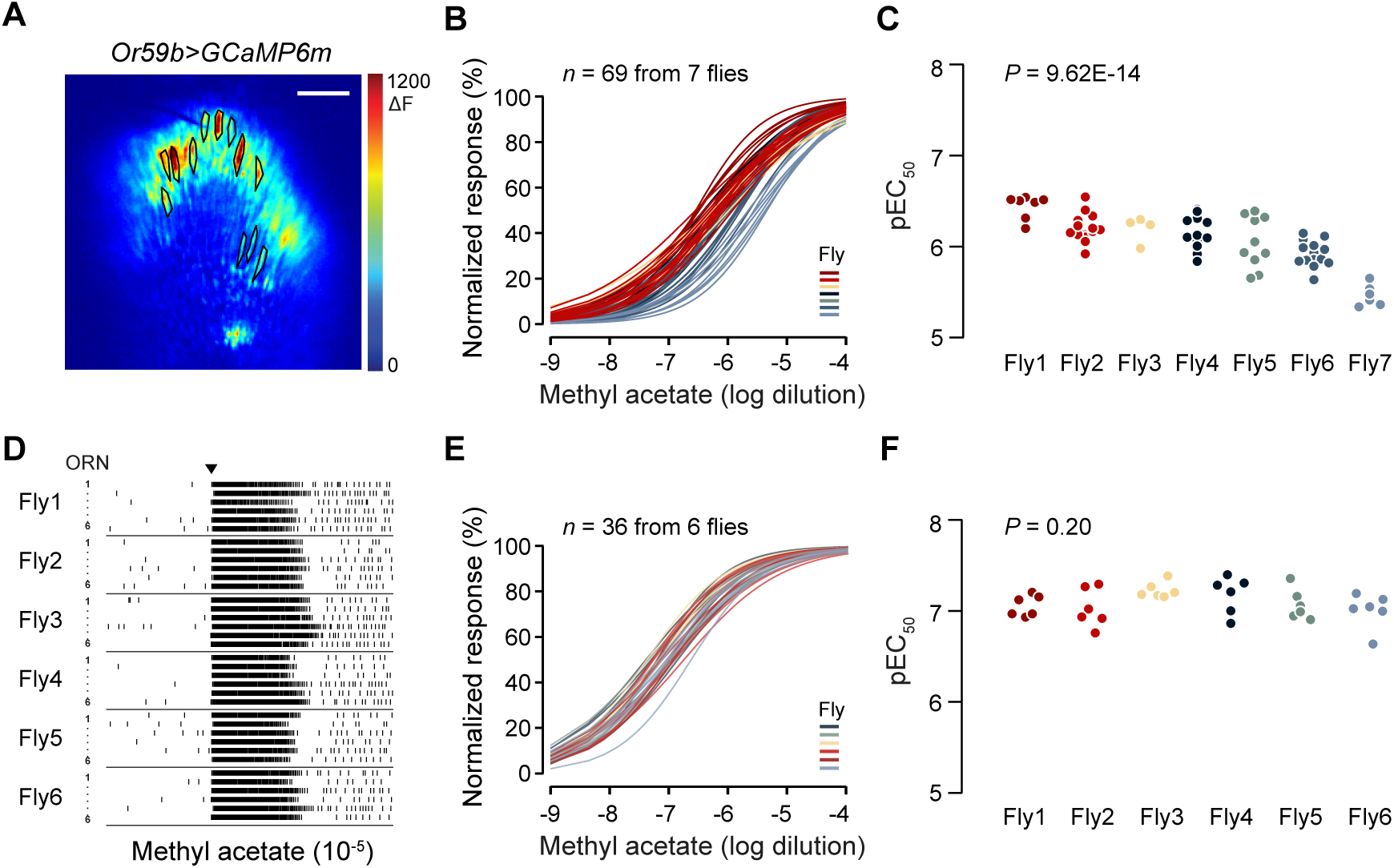
Homotypic ab2A neurons exhibit similar odorant sensitivity. **A.** Odor-evoked calcium fluorescence at the dendrites of ab2A ORNs expressing GCaMP6m (*Or59b>GCaMP6m*). Representative pseudocolored heatmap shows peak ΔF response. Black circles indicate ROIs corresponding to individual GCaMP6m-positive ab2 sensilla. Odor stimulus: methyl acetate (10^-5^). Scale bar: 15 μm. **B.** Dose–response curves of peak calcium responses (ΔF/F), normalized to each neuron’s maximum response to facilitate direct comparison of sensitivity. Each curve represents responses from a single ab2A ORN, fitted with the Hill equation (*n* = 69 ab2 sensilla from 7 flies; mean ± SEM: 9.86 ± 1.53 ab2 sensilla per antenna). Responses from neurons within the same antenna are indicated by the same color. **C.** Quantification of pEC_50_ values from **B**, defined as –logEC_50_. Statistical comparisons between pEC_50_ across flies were performed using one-way ANOVA. **D.** Single-sensillum recordings. Raster plots showing ab2A spike responses. Six ab2A ORNs were recorded per antenna from six flies. Genotype: *Or59b>GCaMP6m* (same as **A**–**C**). Odor stimulus: methyl acetate (10^-5^), onset marked by inverted triangle. **E.** Dose–response curves of peak spike responses, normalized to each neuron’s maximum response. Each curve represents responses from a single ab2A ORN, fitted with the Hill equation (*n* = 36 neurons from 6 flies). Responses recorded from neurons within the same antenna are indicated by the same color. **F.** Quantification of pEC_50_ values from **E**, defined as –logEC_50_. Statistical comparisons between pEC_50_ across flies were performed using one-way ANOVA.

**Figure 2.**
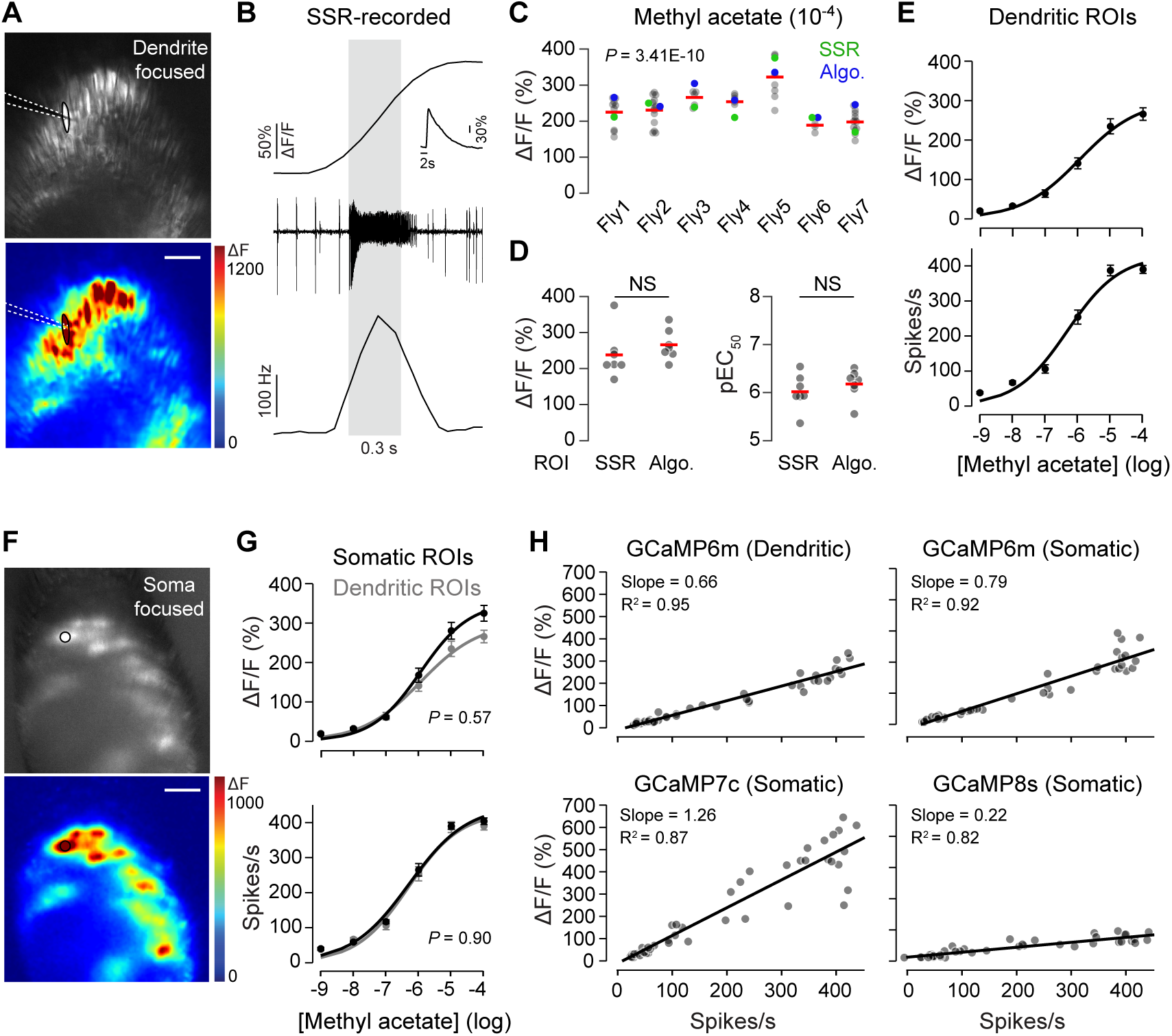
Simultaneous single-sensillum recording and calcium imaging. **A.** Representative raw fluorescence (top) and corresponding pseudocolored heatmap (bottom) showing calcium responses (ΔF) in ab2A neurons from *Or59b>GCaMP6m* flies. The focal plane was positioned at the level of the ab2 sensilla. The recording electrode used for single-sensillum recording (SSR) is outlined with a white dashed line. Scale bar: 15 μm. **B.** Simultaneously recorded calcium and spike responses from the ab2A ORN outlined in **A**. Shown are the calcium fluorescence trace (ΔF/F), the corresponding spike trace, and a peri-stimulus time histogram (PSTH). Gray box indicates a 300-ms methyl acetate stimulus (10^-6^ dilution). Inset: calcium fluorescence trace over a longer duration (22 seconds). **C.** Scatter plot of peak calcium responses (ΔF/F) to methyl acetate (10^-4^ dilution) from individual sensillum ROIs within the same antenna. Each point represents the peak response from one sensillum. The green dot indicates the SSR-recorded sensillum; the blue dot marks the sensillum selected by the analysis algorithm (see Methods). Horizontal bar indicates the mean. Statistical comparisons across flies were performed using one-way ANOVA. **D.** Comparison of peak ΔF/F (left) and pEC₅₀ values (right; defined as –logEC₅₀) between SSR-recorded and algorithm-selected ROIs (*n* = 7 neurons from 7 flies). Statistical test: two-sided Wilcoxon rank-sum test (ΔF/F: *P* = 0.17, pEC_50_: *P* = 0.34). **E.** Dose–response curves of peak calcium signals (top) and peak spike responses (bottom), fitted with the Hill equation. *n* = 7 neurons, with one ab2A neuron per fly. **F**–**G**. As in **A** and **E**, but with the imaging focal plane positioned at the level of the ab2 somas. The algorithm-selected ROI is outlined in black. For comparison, dendritic dose–response curves from panel **E** are shown in gray. Statistical comparison of dendritic vs. somatic dosage curves was performed using two-way ANCOVA. **H.** Linear regression analysis of simultaneously recorded peak calcium and spike responses across odorant dilutions. Each data point represents one odorant concentration from one fly (*n* = 42; 6 dilutions × 7 flies). The slope of best-fit line and coefficient of determination (R²) are shown. Statistical comparisons of spike and somatic calcium response relationships across GCaMP variants were performed using one-way MANOVA: 6m vs. 7c, *P* = 0.32; 6m vs. 8s, *P* = 0.19; 7c vs. 8s, *P* = 0.0042.

## Results

### Homotypic ORNs display similar odor sensitivity

Before establishing the dual recording method, we first examined whether *Drosophila* ORNs that express the same receptor—referred to as homotypic ORNs—share the same odor sensitivity range. Addressing this question is essential for determining how to select regions of interest (ROIs) when analyzing GCaMP fluorescence signals. It is important to understand whether *Drosophila* ORNs, like those in mice, exhibit substantial variability in sensitivity among homotypic ORNs, which can span two to three orders of magnitude (Bozza et al., 2002; Grosmaitre et al., 2006). If such variability exists, it would be necessary to analyze calcium signals from as many ORNs as possible within the field of view to capture the full sensitivity range. In that case, averaging responses across multiple ORNs may obscure meaningful differences and fail to represent the true sensitivity range of the homotypic population. Conversely, if homotypic ORNs exhibit similar sensitivity, recording from a single neuron would be sufficient to represent the entire population.

To address this question, we recorded calcium fluorescence signals in the large spike “A” neuron of the <u>a</u>ntennal <u>b</u>asiconic sensillum type <u>2</u> (ab2A), which expresses the Or59b receptor (Hallem et al., 2004). Calcium signals were visualized by expressing GCaMP6m (Chen et al., 2013) under the control of the *Or59b-GAL4* driver and imaging through the cuticle with widefield epifluorescence microscopy, as described by us and others (Kamikouchi et al., 2010; Knecht et al., 2017; Mariette et al., 2023; Verschut et al., 2023; Vulpe et al., 2021). We focused on ab2A ORNs because this neuronal type exhibits highly elaborate dendritic arborization with over 300 branches filling the entire sensillum lumen (Nava Gonzales et al., 2021). This morphological feature facilitates visualization of dendritic calcium fluorescence by allowing the imaging focal plane to be placed directly at the level of the sensillum cuticle. As a result, we were able to unequivocally define the ROI on the dendrites of each GCaMP-expressing ab2A neuron (Figure 1A). Dendritic ROIs provide a clear and distinct representation of individual ORNs in widefield imaging.

Each *Drosophila* antenna contains approximately 28 ab2 sensilla (Takagi et al., 2024). Within the imaging focal plane, we identified ∼35% of these sensilla per antenna (9.86 ± 1.53, mean ± SEM, *n* = 7 flies). To compare the odorant sensitivity of homotypic ab2A ORNs, we analyzed normalized dose–response curves from all identified dendritic ROIs and calculated their effective concentration 50% (EC_50_) values. The results revealed modest variability, with EC_50_ values spanning approximately 16-fold across all ORNs (Figures 1B and 1C). One-way ANOVA of the pEC_50_ values (–logEC_50_) indicated significant differences across flies (*P* < 0.001), with 70% of the variance attributable to differences between individual flies (Figure 1C). However, within each antenna, the spread of EC_50_ values among homotypic ORNs was remarkably small, averaging only 0.68-fold difference (see Source Data for Figure 1). These results suggest that, unlike in mice, homotypic ORNs in a single *Drosophila* antenna have highly similar sensitivity. Thus, the calcium fluorescence response of a single *Drosophila* ORN is representative of its homotypic population within the same antenna.

What then accounts for the differences in odorant sensitivity across flies observed in the calcium imaging experiments (Figure 1C)? Do these variations reflect true biological heterogeneity between individuals, or do they arise from variability in GCaMP expression? To address this question, we performed single-sensillum recordings (SSR) from multiple homotypic ORNs in flies of the same genotype used in the calcium imaging experiments (*Or59b-GAL4>UAS-GCaMP6m*). This approach allowed us to determine the ground-truth odorant sensitivity of homotypic ORNs both within individual antennae and across flies.

We recorded from six ab2A ORNs per antenna—the maximum number of ab2 sensilla that we could reliably record in every antenna—in a total of six flies. Spike analysis revealed consistent odor-evoked responses across ab2A neurons (Figure 1D). Dose–response curves and EC_50_ values showed a narrow range of sensitivity spread, with EC_50_ values spanning less than 6-fold across all SSR-recorded ORNs (Figures 1E and 1F). One-way ANOVA of the pEC_50_ values revealed no significant differences across flies (*P* = 0.20, Figure 1F), and the average spread of EC_50_ values among homotypic ORNs was also small (0.39-fold; see Source Data for Figure 1). These results support the conclusion that homotypic ab2A ORNs exhibit highly consistent odorant sensitivity and spike response frequencies. They also suggest that the broader EC_50_ variation observed in calcium imaging likely results from variability in GCaMP expression across flies. Together, these results demonstrate that homotypic ORNs in *Drosophila* show uniform odorant sensitivity both within a single antenna and across individual flies, and that the spike response of a single ORN reliably reflects that of its homotypic population.

To further compare the two approaches, we analyzed ab2A sensitivities as measured by calcium imaging and SSR, respectively (Figure 1). pEC_50_ values were consistently lower with calcium imaging (mean ± SEM: 6.07 ± 0.04) compared to SSR (7.09 ± 0.03), reflecting an approximately 10-fold difference in apparent odorant sensitivity (Mann–Whitney rank-sum test, *P* < 0.001).

This discrepancy likely reflects the higher sensitivity of SSR in detecting neuronal responses. Notably, although the measured absolute sensitivity differed between calcium imaging and SSR, the spread of pEC_50_ values did not differ significantly between the two methods (*P* = 0.836). This result indicates that widefield calcium imaging—despite potential variability in ROI selection or focal plane—does not introduce artifactual variability and provides a reliable measure of ORN sensitivity across the population.

### Dual recording: simultaneous calcium imaging and single-sensillum recording

Having established the similarity of odorant sensitivity across homotypic ORNs, we next optimized a dual-modality recording method. In brief, we simultaneously recorded GCaMP fluorescence signals from ab2A neurons using widefield imaging of the antenna, and recorded their spike activity via single-sensillum recordings in response to airborne odor stimulation. This combined approach allows concurrent acquisition of both calcium and electrophysiological responses from the same antenna.

We first performed dual recordings with the imaging focal plane positioned at the level of the sensillum cuticle, allowing us to place ROIs on all sensilla containing GCaMP-expressing ab2A, including the one targeted for SSR (Figure 2A). This setup enabled direct comparison of calcium and spike responses from the same neuron, rather than correlating data across different flies and separate experiments, as conducted in previous studies (Mariette et al., 2023). While spike activity closely followed the onset and offset of the odor stimulus with rapid rise and decay (τ_on_ = 244 ms and τ_off_ = 274 ms), the calcium fluorescence signal displayed considerably slower dynamics (τ_on_ = 485 ms and τ_off_ = 3920 ms, Figure 2B). As a result, the peaks of the spike and calcium responses were not temporally aligned. Given this temporal mismatch, our subsequent analysis of peak calcium and corresponding spike responses was not intended to define a precise transfer function between action potentials and GCaMP fluorescence. Rather, our goal was to provide a reference framework for interpreting calcium response amplitudes in individual ORN types—for example, to relate a given peak ΔF/F value to an approximate peak firing rate.

Given that the calcium and spike responses of a single ORN can represent those of its homotypic counterparts within the same antenna (Figure 1), is it necessary to correlate these two responses from the same neuron to accurately assess their relationship? To address this question, we compared the calcium response from the SSR-recorded ROI with those from all other ROIs in the same antenna, including the one identified by our analysis algorithm as showing the strongest response to the odorant methyl acetate (Figure 2C; see Methods for details). Consistent with our earlier survey using widefield calcium imaging (Figures 1B and 1C), we observed significant variability in methyl acetate-evoked calcium responses across flies (Figure 2C). However, statistical analysis showed no significant difference in either fluorescence signals (ΔF/F) or EC_50_ between the SSR-recorded and algorithm-selected ROIs (Figure 2D).

These results suggest that, for the purposes of correlation analysis, it is not strictly necessary to match calcium and spike recordings from the exact same neuron. Thus, our subsequent correlation analyses remain valid even when the algorithm-selected ROI does not always correspond to the SSR-recorded neuron.

This consideration is particularly relevant for future studies seeking to extend correlation analysis to other ORN types—such as trichoid, coeloconic, or small-spike basiconic neurons— which possess relatively few dendritic branches (Nava Gonzales et al., 2021). As such, their dendritic GCaMP fluorescence may be too weak to detect under widefield imaging, making it challenging to identify ROIs corresponding to SSR-recorded sensilla. To facilitate such future large-scale investigations, we chose to use algorithm-selected ROIs to define calcium dose– response relationships that can be correlated with spike dose–response curves from homotypic ORNs within the same antenna (Figure 2E).

To broaden the utility of dual recordings, we next compared dendritic and somatic calcium responses in algorithm-selected ROIs from *Or59b-GAL4>UAS-GCaMP6m* flies, given that only a limited subset of large basiconic ORNs are amenable to dendritic imaging under widefield microscopy. To this end, we conducted additional dual recordings while focusing the imaging plane on the ORN somas (Figure 2F). Comparison of the dendritic and somatic calcium dose– response curves revealed no significant difference between the two focal planes (Figure 2G, top panel). As expected, the corresponding spike dose–response curves were virtually identical between these two conditions (Figure 2G, bottom panel, black and gray curves). Collectively, these results support the validity of using somatic calcium signals from algorithm-selected ROIs to correlate with spike responses.

To determine the relationship between calcium and spike responses in ab2A ORNs, we plotted the peak calcium fluorescence signals (ΔF/F, %) against the peak spike responses (Hz). Remarkably, a strong linear correlation was observed across the full range of spike firing frequencies, spanning over 400 Hz. This relationship remained consistent regardless of the calcium signal source (dendrites or soma) or the specific GCaMP indicators used (Figure 2H). Notably, while the linear relationship was maintained in two other GCaMP variants (Dana et al., 2019; Zhang et al., 2023), the slopes differed depending on the indicator. GCaMP7c exhibited a similar spike–calcium response relationship to GCaMP6m, whereas GCaMP8s showed a significantly shallower slope compared to GCaMP7c (MANOVA followed by Tukey–Kramer test: 6m vs. 7c, *P* = 0.32; 6m vs. 8s, *P* = 0.19; 7c vs. 8s, *P* = 0.0042; Figure 2H).

### Linear relationship between calcium and spike responses is odorant-invariant

Is the relationship of the calcium–spike correlation affected by odorant identity? This question is important because most tuning odorant receptors in *D. melanogaster* respond to multiple odorants with varying sensitivity (Hallem & Carlson, 2006). To address this question, we performed dual recordings in ab2A ORNs of *Or59b-GAL4>UAS-GCaMP6m* flies using another odorant, ethyl acetate, which also strongly activates the Or59b receptor but is less effective than methyl acetate (Hallem & Carlson, 2006; Zhang et al., 2019). Notably, the linear relationship between calcium and spike responses with ethyl acetate was virtually identical to that obtained with methyl acetate (one-way MANOVA, *P* = 0.36; Figure 3A).

**Figure 3.**
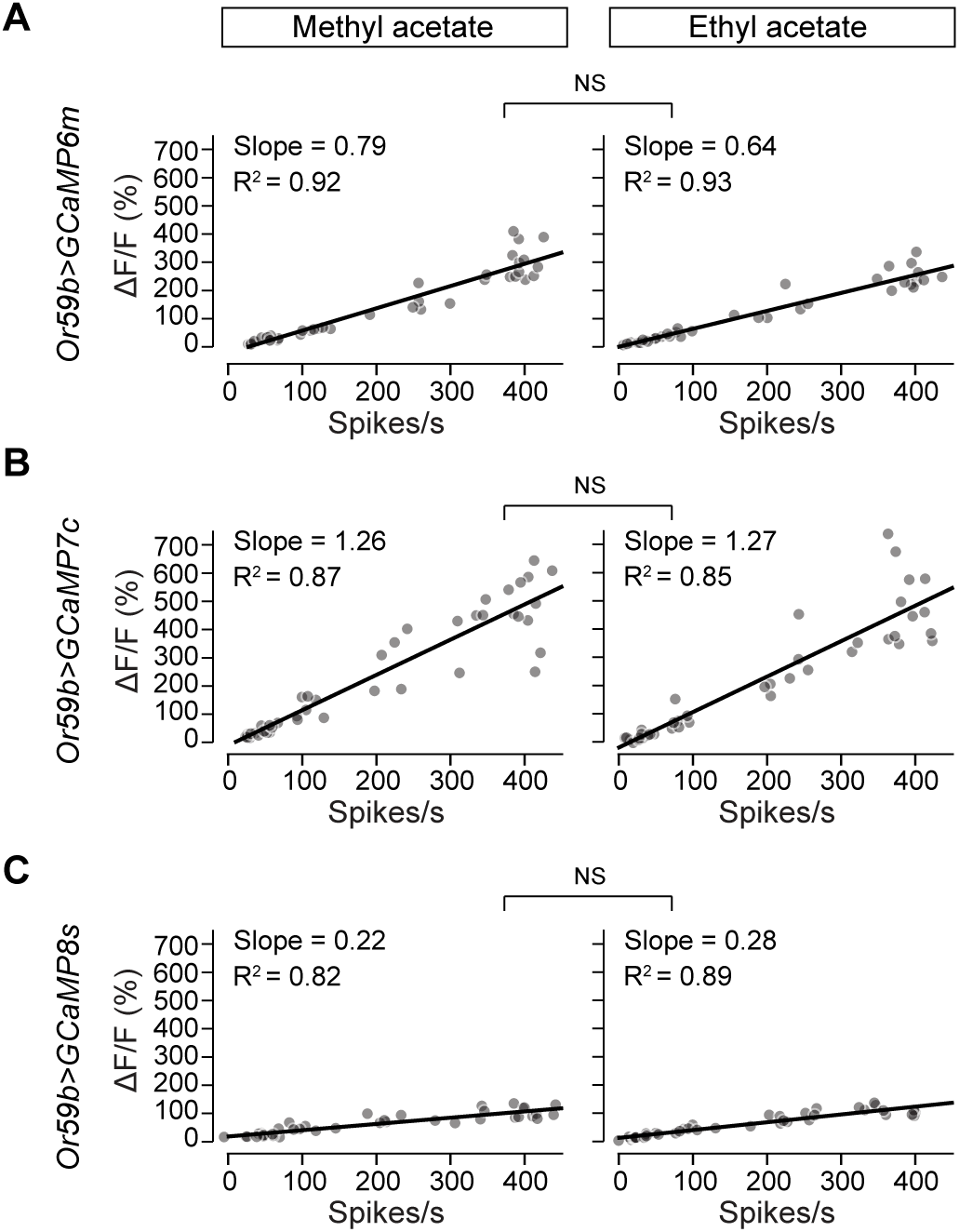
Linear relationship between calcium and spike responses is odorant-invariant. **A.** Linear regression analysis of simultaneously recorded peak calcium and spike responses across different concentrations of odor stimuli in ab2A ORNs expressing GCaMP6m (*Or59b>GCaMP6m*). Left: responses to methyl acetate (same data as shown in Figure 2H). Right: responses to ethyl acetate. Each data point represents one odorant concentration from one fly (*n* = 36–42; 6 dilutions × 6–7 flies). The slope of best-fit line and coefficient of determination (R²) are shown. Statistical comparisons between responses of different odorants were performed using one-way MANOVA (*P* = 0.36). **B**–**C**. Same as in A, except that recordings were performed in ab2A ORNs expressing GCaMP7c (**B**) or GCaMP8s (**C**). *P* = 0.80 for GCaMP7c and *P* = 0.27 for GCaMP8s.

We extended this analysis to two additional GCaMP variants, GCaMP7c and GCaMP8s (Dana et al., 2019; Zhang et al., 2023). As observed previously, GCaMP8s exhibited a shallower slope compared to GCaMP7c (Figure 2H), but within each variant, the relationship remained unchanged between methyl acetate and ethyl acetate (Figures 3B and 3C). Together, these results indicate that the linear relationship between calcium and spike responses in *Drosophila* ORNs is an intrinsic property of the neurons and is invariant to odorant identity.

### Linear relationship between calcium and spike responses is preserved across ages

Thus far, we examined *Or59b-GAL4>UAS-GCaMP* flies that were four days post-eclosion. Given that GAL4-driven UAS expression tends to increase with age in adult flies, especially beyond 5 days (Seroude et al., 2002), we extended our analysis to *Or59b-GAL4>UAS-GCaMP6m* flies that were three, five, and seven days old.

We first compared baseline fluorescence levels (F_0_ – F_bac_, see Methods for details), which reflect basal calcium levels, GCaMP6m expression, or a combination of both. Baseline fluorescence in ab2A ORNs did not differ significantly between 3-day- and 5-day-old flies. Unexpectedly, fluorescence level was significantly lower in 7-day-old flies compared to 5-day-olds (Figure 4A), suggesting that basal GCaMP signal does not necessarily increase with age in transgenic flies.

**Figure 4.**
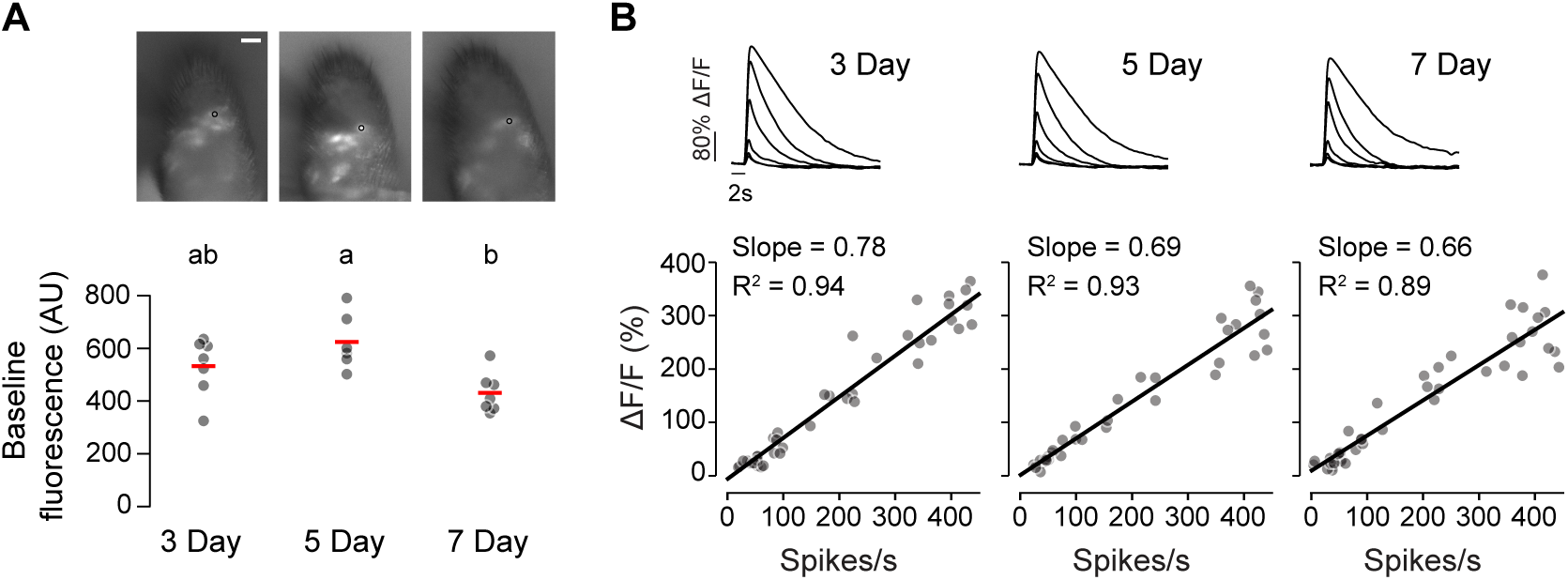
Linear relationship between calcium and spike responses is preserved across ages. **A.** Baseline fluorescence of ab2A ORNs expressing GCaMP6m (*Or59b>GCaMP6m*) in flies at 3, 5, and 7 days post-eclosion. Top: representative raw fluorescence image. Scale bar: 15 µm. Bottom: quantification of baseline fluorescence intensity across the three age groups. Each data point represents the pixel value from an algorithm-selected ROI per antenna. Horizontal bars indicate group means. Statistical comparisons were performed using one-way ANOVA followed by the Tukey–Kramer test (*n* = 6–7 antenna per age group). Groups with significant differences (*P* < 0.05) are labeled with different letters. **B.** Linear regression analysis of simultaneously recorded peak calcium and spike responses to methyl acetate. Top: average GCaMP fluorescence transients in response to increasing concentrations of methyl acetate (stimulus duration: 300 ms). Bottom: Liner regression analysis. Each data point represents one odorant concentration from one fly (*n* = 36–42; 6 dilutions × 6–7 flies). The slope of best-fit line and coefficient of determination (R²) are shown. One-way MANOVA: *P* > 0.05 across age groups.

Dual recordings revealed that the linear relationship between calcium and spike responses was preserved across all three age groups. The relationships were also similar (one-way MANOVA: 3d vs. 5d, *P* = 0.96; 3d vs. 7d, *P* = 0.94; 5d vs. 7d, *P* = 1.00; Figure 4B), indicating that age-related variation in GCaMP expression does not affect the fidelity of spike–calcium relationship in ab2A ORNs. These results support the robustness of the spike–calcium relationship in *Drosophila* ORNs across a range of early adult ages.

## Discussion

In this study, we established a robust method for simultaneously recording odor-evoked spike activity and calcium responses from ORNs in *D. melanogaster*. By directly comparing these two readouts across different imaging planes, GCaMP variants, odorants, and fly ages, we found that the linear relationship between spike firing and calcium fluorescence is remarkably stable. While the slope of this relationship can vary depending on the specific GCaMP indicator used, the linearity itself remains consistent. This consistency supports the use of calcium imaging as a reliable proxy for neuronal activity in *Drosophila* ORNs, thus facilitating its broader application in functional studies of olfactory coding. Importantly, this robust relationship across multiple conditions forms the foundation for future large-scale surveys of spike–calcium response relationship across diverse ORN types.

Our results also show that homotypic ORNs in *Drosophila*, such as ab2A neurons expressing the Or59b receptor, exhibit similar odor sensitivity within the same antenna and across flies. Despite previously documented morphological variability among homotypic neurons—for example, differences in dendritic size and morphology observed in the ab1C or ab1D population (Choy et al., 2025)—this possible structural heterogeneity does not appear to result in functional variability in odor sensitivity. In contrast to the wide range of sensitivities seen in rodent ORNs expressing the same receptor (Bozza et al., 2002; Grosmaitre et al., 2006), the similarity among homotypic *Drosophila* ORNs greatly simplifies analysis. It also validates the use of single-sensillum recordings or single-cell imaging as representative of the entire homotypic population, thereby enabling more efficient and scalable characterization of odor coding at the level of individual ORN types.

## Supporting information

Source Data for Figure 1

Source Data for Figure 2

Source Data for Figure 3

Source Data for Figure 4

## Acknowledgments

We thank Carlotta Martelli for sharing the MATLAB code for spike analysis, and Renny Ng for comments on the manuscript.

## Disclosure statement

No potential conflict of interest was reported by the authors.

## Author contributions

Y. Xiao performed dual recordings and data analysis, and developed data analysis code in MATLAB. S-T.W. optimized the dual recording method with help from C-Y.S., taught Y. Xiao calcium imaging, and helped with fly work. Y. Xuan developed the initial data analysis code in Python. S.A.R. assisted with statistical analysis. C-Y.S. conceived the project, acquired funding, performed single-sensillum recordings for Figure 2, and supervised the project. Y. Xiao and C-Y.S. wrote the manuscript with inputs from all authors.

## Funding

This work was supported by NIH grants to C-Y.S. (R21DC020536, R21AI169343, R01DC016466, and R01DC021551).

## Data availability statement

All fly lines used in this study are available at the Bloomington *Drosophila* Stock Center. For additional information or data requests, please contact the corresponding author, Chih-Ying Su.

## Notes

### Competing Interest Statement

The authors have declared no competing interest.

## References

Afify, A., Betz, J. F., Riabinina, O., Lahondère, C., & Potter, C. J. (2019). Commonly Used Insect Repellents Hide Human Odors from Anopheles Mosquitoes. Current Biology, 29(21), 3669–3680.e5. 10.1016/j.cub.2019.09.007

Ali, F., & Kwan, A. C. (2019). Interpreting in vivo calcium signals from neuronal cell bodies, axons, and dendrites: a review. Neurophotonics, 7(01), 1. 10.1117/1.nph.7.1.011402

Bonheur, M., Swartz, K. J., Metcalf, M. G., Wen, X., Zhukovskaya, A., Mehta, A., Connors, K. E., Barasch, J. G., Jamieson, A. R., Martin, K. C., Axel, R., & Hattori, D. (2023). A rapid and bidirectional reporter of neural activity reveals neural correlates of social behaviors in Drosophila. Nature Neuroscience, 26(7), 1295–1307. 10.1038/s41593-023-01357-w

Bozza, T., Feinstein, P., Zheng, C., & Mombaerts, P. (2002). Odorant receptor expression defines functional units in the mouse olfactory system. Journal of Neuroscience, 22(8), 3033–3043. 10.1523/jneurosci.22-08-03033.2002

Bui, M., Shyong, J., Lutz, E. K., Yang, T., Li, M., Truong, K., Arvidson, R., Buchman, A., Riffell, J. A., & Akbari, O. S. (2019). Live calcium imaging of Aedes aegypti neuronal tissues reveals differential importance of chemosensory systems for life-history-specific foraging strategies. BMC Neuroscience, 20(1), 27. 10.1186/s12868-019-0511-y

Cao, S., Huang, T., Shen, J., Liu, Y., & Wang, G. (2020). An Orphan Pheromone Receptor Affects the Mating Behavior of Helicoverpa armigera. Frontiers in Physiology, 11(April), 1–10. 10.3389/fphys.2020.00413

Carcaud, J., Otte, M., Grünewald, B., Haase, A., Sandoz, J. C., & Beye, M. (2023). Multisite imaging of neural activity using a genetically encoded calcium sensor in the honey bee. PLoS Biology, 21(1). 10.1371/journal.pbio.3001984

Chang, H., Liu, Y., Ai, D., Jiang, X., Dong, S., & Wang, G. (2017). A Pheromone Antagonist Regulates Optimal Mating Time in the Moth Helicoverpa armigera. Current Biology, 27(11), 1610–1615.e3. 10.1016/j.cub.2017.04.035

Chen, T.-W., Wardill, T. J., Sun, Y., Pulver, S. R., Renninger, S. L., Baohan, A., Schreiter, E. R., Kerr, R. a., Orger, M. B., Jayaraman, V., Looger, L. L., Svoboda, K., & Kim, D. S. (2013). Ultrasensitive fluorescent proteins for imaging neuronal activity. Nature, 499(7458), 295–300. 10.1038/nature12354

Choy, J., Charara, S., Cauwenberghs, K., McKaughan, Q., Kim, K., Ellisman, M. H., & Su, C.-Y. (2025). Population-level morphological analysis of paired CO2- and odor-sensing olfactory neurons in D. melanogaster via volume electron microscopy. In eLife (p. RP106389). 10.7554/eLife.106389.1

Dana, H., Sun, Y., Mohar, B., Hulse, B. K., Kerlin, A. M., Hasseman, J. P., Tsegaye, G., Tsang, A., Wong, A., Patel, R., Macklin, J. J., Chen, Y., Konnerth, A., Jayaraman, V., Looger, L. L., Schreiter, E. R., Svoboda, K., & Kim, D. S. (2019). High-performance calcium sensors for imaging activity in neuronal populations and microcompartments. Nature Methods, 16(7), 649–657. 10.1038/s41592-019-0435-6

DeGennaro, M., McBride, C. S., Seeholzer, L., Nakagawa, T., Dennis, E. J., Goldman, C., Jasinskiene, N., James, A. A., & Vosshall, L. B. (2013). orco mutant mosquitoes lose strong preference for humans and are not repelled by volatile DEET. Nature, 498(7455), 487–491. 10.1038/nature12206

Dormont, L., Mulatier, M., Carrasco, D., & Cohuet, A. (2021). Mosquito Attractants. Journal of Chemical Ecology, 47(4–5), 351–393. 10.1007/s10886-021-01261-2

Dubbs, A., Guevara, J., & Yuste, R. (2016). moco: Fast motion correction for calcium imaging. Frontiers in Neuroinformatics, 10(FEB), 1–4. 10.3389/fninf.2016.00006

Edelstein, A. D., Tsuchida, M. A., Amodaj, N., Pinkard, H., Vale, R. D., & Stuurman, N. (2014). Advanced methods of microscope control using μManager software. Journal of Biological Methods, 1(2), e10. 10.14440/jbm.2014.36

Fandino, R. A., Haverkamp, A., Bisch-Knaden, S., Zhang, J., Bucks, S., Nguyen, T. A. T., Schröder, K., Werckenthin, A., Rybak, J., Stengl, M., Knaden, M., Hansson, B. S., & Große-Wilde, E. (2019). Mutagenesis of odorant coreceptor Orco fully disrupts foraging but not oviposition behaviors in the hawkmoth Manduca sexta. Proceedings of the National Academy of Sciences, 116(31), 15677–15685. 10.1073/pnas.1902089116

Fosque, B. F., Sun, Y., Dana, H., Yang, C. T., Ohyama, T., Tadross, M. R., Patel, R., Zlatic, M., Kim, D. S., Ahrens, M. B., Jayaraman, V., Looger, L. L., & Schreiter, E. R. (2015). Labeling of active neural circuits in vivo with designed calcium integrators. Science, 347(6223), 755–760. 10.1126/science.1260922

Friel, D. D., & Chiel, H. J. (2008). Calcium dynamics: analyzing the Ca2+ regulatory network in intact cells. Trends in Neurosciences, 31(1), 8–19. 10.1016/j.tins.2007.11.004

Fujiwara, T., Kazawa, T., Sakurai, T., Fukushima, R., Uchino, K., Yamagata, T., Namiki, S., Haupt, S. S., & Kanzaki, R. (2014). Odorant concentration differentiator for intermittent olfactory signals. Journal of Neuroscience, 34(50), 16581–16593. 10.1523/JNEUROSCI.2319-14.2014

Gao, X. J., Riabinina, O., Li, J., Potter, C. J., Clandinin, T. R., & Luo, L. (2015). A transcriptional reporter of intracellular Ca2+ in Drosophila. Nature Neuroscience, 18(6), 917–925. 10.1038/nn.4016

Grabe, V., Baschwitz, A., Dweck, H. K. M., Lavista-Llanos, S., Hansson, B. S., & Sachse, S. (2016). Elucidating the neuronal architecture of olfactory glomeruli in the Drosophila antennal lobe. Cell Reports, 16(12), 3401–3413. 10.1016/j.celrep.2016.08.063

Grosmaitre, X., Vassalli, A., Mombaerts, P., Shepherd, G. M., & Ma, M. (2006). Odorant of olfactory neurons expressing the odorant receptor MOR23: A patch clamp analysis in gene-targeted mice. Proceedings of the National Academy of Sciences of the United States of America, 103(6), 1970–1975. 10.1073/pnas.0508491103

Guo, X., Yu, Q., Chen, D., Wei, J., Yang, P., Yu, J., Wang, X., & Kang, L. (2020). 4-Vinylanisole is an aggregation pheromone in locusts. Nature, February. 10.1038/s41586-020-2610-4

Hallem, E. A., & Carlson, J. R. (2006). Coding of Odors by a Receptor Repertoire. Cell, 125(1), 143–160. 10.1016/j.cell.2006.01.050

Hallem, E. A., Ho, M. G., & Carlson, J. R. (2004). The Molecular Basis of Odor Coding in the Drosophila Antenna. Cell, 117(7), 965–979. 10.1016/j.cell.2004.05.012

Hart, T., Frank, D. D., Lopes, L. E., Olivos-Cisneros, L., Lacy, K. D., Trible, W., Ritger, A., Valdés-Rodríguez, S., & Kronauer, D. J. C. (2023). Sparse and stereotyped encoding implicates a core glomerulus for ant alarm behavior. Cell, 186(14), 3079–3094.e17. 10.1016/j.cell.2023.05.025

Helmchen, F., & Tank, D. W. (2015). A single-compartment model of calcium dynamics in nerve terminals and dendrites. Cold Spring Harbor Protocols, 2015(2), 155–167. 10.1101/pdb.top085910

Huang, L., Ledochowitsch, P., Knoblich, U., Lecoq, J., Murphy, G. J., Reid, R. C., de Vries, S. E. J., Koch, C., Zeng, H., Buice, M. A., Waters, J., & Li, L. (2021). Relationship between simultaneously recorded spiking activity and fluorescence signal in gcamp6 transgenic mice. ELife, 10, 1–19. 10.7554/eLife.51675

Jiang, X., Dimitriou, E., Grabe, V., Sun, R., Chang, H., Zhang, Y., Gershenzon, J., Rybak, J., Hansson, B. S., & Sachse, S. (2024). Ring-shaped odor coding in the antennal lobe of migratory locusts. Cell, 3973–3991. 10.1016/j.cell.2024.05.036

Kamikouchi, A., Wiek, R., Effertz, T., Göpfert, M. C., & Fiala, A. (2010). Transcuticular optical imaging of stimulus-evoked neural activities in the Drosophila peripheral nervous system. Nature Protocols, 5(7), 1229–1235. 10.1038/nprot.2010.85

Kim, S. M., Su, C. Y., & Wang, J. W. (2017). Neuromodulation of Innate Behaviors in Drosophila. Annual Review of Neuroscience, 40(1), 327–348. 10.1146/annurev-neuro-072116-031558

Knecht, Z. A., Silbering, A. F., Cruz, J., Yang, L., Croset, V., Benton, R., & Garrity, P. A. (2017). Ionotropic receptor-dependent moist and dry cells control hygrosensation in Drosophila. ELife, 6, e26654. 10.7554/eLife.26654

Konopka, J. K., Task, D., Afify, A., Raji, J., Deibel, K., Maguire, S., Lawrence, R., & Potter, C. J. (2021). Olfaction in Anopheles mosquitoes. Chemical Senses, 46, 1–10. 10.1093/chemse/bjab021

Lahondère, C., Vinauger, C., Okubo, R. P., Wolff, G. H., Chan, J. K., Akbari, O. S., & Riffell, J. A. (2020). The olfactory basis of orchid pollination by mosquitoes. Proceedings of the National Academy of Sciences of the United States of America, 117(1), 708–716. 10.1073/pnas.1910589117

Liu, X., Zhang, J., Yan, Q., Miao, C., Han, W., Hou, W., Yang, K., Hansson, B. S., Peng, Y.-C., Guo, J.-M., Xu, H., Wang, C., Dong, S., & Knaden, M. (2020). The Molecular Basis of Host Selection in a Crucifer-Specialized Moth. Current Biology, 30(22), 4476–4482.e5. 10.1016/j.cub.2020.08.047

Manwill, P. K., Kalsi, M., Wu, S., Martinez-Rodriguez, E. J., Cheng, X., Piermarini, P. M., & Rakotondraibe, H. L. (2020). Semi-synthetic cinnamodial analogues: Structural insights into the insecticidal and antifeedant activities of drimane sesquiterpenes against the mosquito Aedes Aegypti. PLoS Neglected Tropical Diseases, 14(2), 1–21. 10.1371/journal.pntd.0008073

Mariette, J., Noël, A., Louis, T., Montagné, N., Chertemps, T., Jacquin-Joly, E., Marion-Poll, F., & Sandoz, J. C. (2023). Transcuticular calcium imaging as a tool for the functional study of insect odorant receptors. Frontiers in Molecular Neuroscience, 16(August), 1–14. 10.3389/fnmol.2023.1182361

Martelli, C., & Fiala, A. (2019). Slow presynaptic mechanisms that mediate adaptation in the olfactory pathway of Drosophila. ELife, 8, 1–26. 10.7554/eLife.43735

Masuyama, K., Zhang, Y., Rao, Y., & Wang, J. W. (2012). Mapping neural circuits with activity-dependent nuclear import of a transcription factor. Journal of Neurogenetics, 26(1), 89–102. 10.3109/01677063.2011.642910

Matthews, B. J., Younger, M. A., & Vosshall, L. B. (2019). The ion channel ppk301 controls freshwater egg-laying in the mosquito aedes aegypti. ELife, 8, 1–27. 10.7554/eLife.43963

McBride, C. S., Baier, F., Omondi, A. B., Spitzer, S. A., Lutomiah, J., Sang, R., Ignell, R., & Vosshall, L. B. (2014). Evolution of mosquito preference for humans linked to an odorant receptor. Nature, 515(7526), 222–227. 10.1038/nature13964

McMeniman, C. J., Corfas, R. A., Matthews, B. J., Ritchie, S. A., & Vosshall, L. B. (2014). Multimodal Integration of Carbon Dioxide and Other Sensory Cues Drives Mosquito Attraction to Humans. Cell, 156(5), 1060–1071. 10.1016/j.cell.2013.12.044

Melo, N., Wolff, G. H., Costa-da-Silva, A. L., Arribas, R., Triana, M. F., Gugger, M., Riffell, J. A., DeGennaro, M., & Stensmyr, M. C. (2019). Geosmin Attracts Aedes aegypti Mosquitoes to Oviposition Sites. Current Biology, 598698. 10.1016/j.cub.2019.11.002

Nava Gonzales, C., McKaughan, Q., Bushong, E. A., Cauwenberghs, K., Ng, R., Madany, M., Ellisman, M. H., & Su, C. Y. (2021). Systematic morphological and morphometric analysis of identified olfactory receptor neurons in drosophila melanogaster. ELife, 10, 2021.04.28.441861. 10.7554/eLife.69896

Ng, R., Wu, S., & Su, C. (2020). Neuronal Compartmentalization: A Means to Integrate Sensory Input at the Earliest Stage of Information Processing? BioEssays, 42(8), 1–11. 10.1002/bies.202000026

Raji, J. I., Melo, N., Castillo, J. S., Gonzalez, S., Saldana, V., Stensmyr, M. C., & DeGennaro, M. (2019). Aedes aegypti Mosquitoes Detect Acidic Volatiles Found in Human Odor Using the IR8a Pathway. Current Biology, 29(8), 1253–1262.e7. 10.1016/j.cub.2019.02.045

Seroude, L., Brummel, T., Kapahi, P., & Benzer, S. (2002). Spatio-temporal analysis of gene expression during aging in Drosophila melanogaster. Aging Cell, 1(1), 47–56. 10.1046/j.1474-9728.2002.00007.x

Shanbhag, S. R., Müller, B., & Steinbrecht, R. A. (1999). Atlas of olfactory organs of Drosophila melanogaster 1. Types, external organization, innervation and distribution of olfactory sensilla. International Journal of Insect Morphology and Embryology, 28(4), 377–397. 10.1016/S0020-7322(99)00039-2

Stewart, B. A., Atwood, H. L., Renger, J. J., Wang, J., & Wu, C.-F. (1994). Improved stability of Drosophila larval neuromuscular preparations in haemolymph-like physiological solutions. Journal of Comparative Physiology A, 175(2), 179–191. 10.1007/BF00215114

Takagi, S., Sancer, G., Abuin, L., Stupski, S. D., Roman Arguello, J., Prieto-Godino, L. L., Stern, D. L., Cruchet, S., Álvarez-Ocaña, R., Wienecke, C. F. R., van Breugel, F., Jeanne, J. M., Auer, T. O., & Benton, R. (2024). Olfactory sensory neuron population expansions influence projection neuron adaptation and enhance odour tracking. Nature Communications, 15(1), 1–18. 10.1038/s41467-024-50808-w

Verschut, T. A., Ng, R., Doubovetzky, N. P., Le Calvez, G., Sneep, J. L., Minnaard, A. J., Su, C. Y., Carlsson, M. A., Wertheim, B., & Billeter, J. C. (2023). Aggregation pheromones have a non-linear effect on oviposition behavior in Drosophila melanogaster. Nature Communications, 14(1), 1544. 10.1038/s41467-023-37046-2

Vinauger, C., Lahondère, C., Wolff, G. H., Locke, L. T., Liaw, J. E., Parrish, J. Z., Akbari, O. S., Dickinson, M. H., & Riffell, J. A. (2018). Modulation of Host Learning in Aedes aegypti Mosquitoes. Current Biology, 28(3), 333–344.e8. 10.1016/j.cub.2017.12.015

Vulpe, A., Kim, H. S., Ballou, S., Wu, S. T., Grabe, V., Nava Gonzales, C., Liang, T., Sachse, S., Jeanne, J. M., Su, C. Y., & Menuz, K. (2021). An ammonium transporter is a non-canonical olfactory receptor for ammonia. Current Biology, 31(15), 3382–3390.e7. 10.1016/j.cub.2021.05.025

Wang, J. W., Wong, A. M., Flores, J., Vosshall, L. B., & Axel, R. (2003). Two-photon calcium imaging reveals an odor-evoked map of activity in the fly brain. Cell, 112(2), 271–282. 10.1016/S0092-8674(03)00004-7

Wheelwright, M., Whittle, C. R., & Riabinina, O. (2021). Olfactory systems across mosquito species. Cell and Tissue Research, 0123456789, 1–16. http://link.springer.com/10.1007/s00441-020-03407-2

Wolff, G. H., Lahondère, C., Vinauger, C., Rylance, E., & Riffell, J. A. (2023). Neuromodulation and differential learning across mosquito species. Proceedings of the Royal Society B: Biological Sciences, 290(1990), 755017. 10.1098/rspb.2022.2118

Zhang, Y., Rózsa, M., Liang, Y., Bushey, D., Wei, Z., Zheng, J., Reep, D., Broussard, G. J., Tsang, A., Tsegaye, G., Narayan, S., Obara, C. J., Lim, J. X., Patel, R., Zhang, R., Ahrens, M. B., Turner, G. C., Wang, S. S. H., Korff, W. L.,… Looger, L. L. (2023). Fast and sensitive GCaMP calcium indicators for imaging neural populations. Nature, 615(7954), 884–891. 10.1038/s41586-023-05828-9

Zhang, Y., Tsang, T. K., Bushong, E. A., Chu, L.-A., Chiang, A.-S., Ellisman, M. H., Reingruber, J., & Su, C.-Y. (2019). Asymmetric ephaptic inhibition between compartmentalized olfactory receptor neurons. Nature Communications, 10(1), 1560. 10.1038/s41467-019-09346-z

Zhao, Z., Zung, J. L., Hinze, A., Kriete, A. L., Iqbal, A., Younger, M. A., Matthews, B. J., Merhof, D., Thiberge, S., Ignell, R., Strauch, M., & McBride, C. S. (2022). Mosquito brains encode unique features of human odour to drive host seeking. Nature, 605(7911), 706–712. 10.1038/s41586-022-04675-4

